## Supplementary Information for "BBSome-dependent ciliary Hedgehog signaling governs cell fate in the white adipose tissue"

### Supplementary Figures

### Figure S1

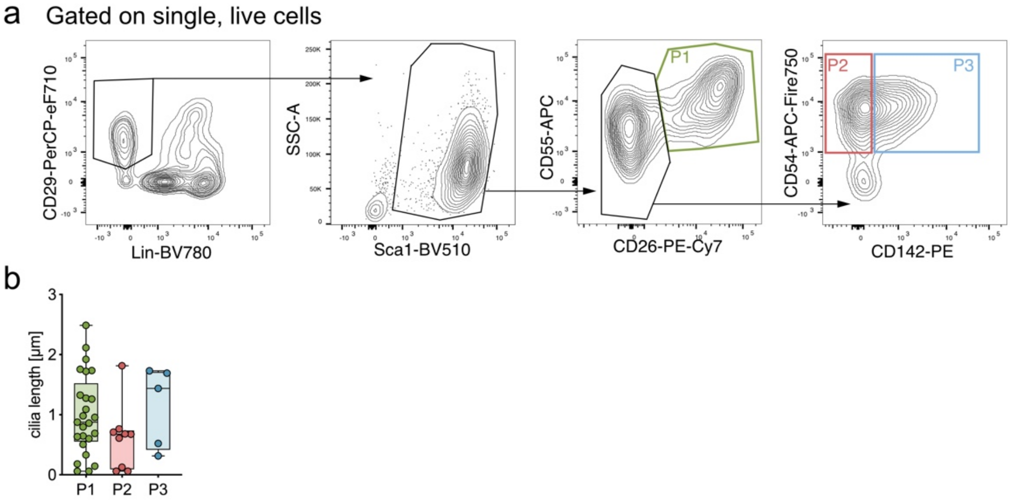


**Fig. S1. Primary cilia regulate the adipogenic potential of adipocytes precursors subpopulations (APCs). a,** Simplified gating strategy to identify the three APC subpopulations using flow cytometry. **b**, Ciliary length of the three subpopulations measured by the ARL13B mask via CiliaQ. (P1 = 25 cilia, P2 = 9 cilia, P3 = 5 cilia, all biological replicates).

### Figure S2

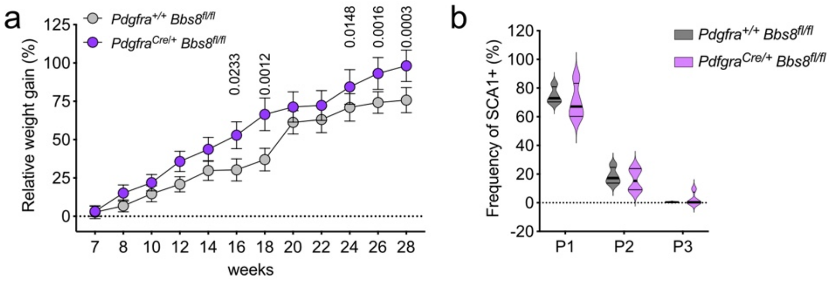


**Fig. S2. Loss of BBS8 in APCs results in obesity. a**, Relative body weight of chow diet-fed *Pdgfra*$^{+/+}$*Bbs8*$^{fl/fl}$ and *Pdgfra*$^{cre/+}$ *Bbs8*$^{fl/fl}$ mice. Weights were normalized to the mean body weight of their respective genotype at the lean time point (7 weeks). Data are shown mean ± SEM, p-values were determined using a Two-way ANOVA with repeated measurements (mixed models). Post-hoc p-value correction for multiple testing was performed using Bonferroni adjustment (n = 3-9). **b,** Frequency distribution of P1-P3 from the total APC pool from *Pdgfra*$^{+/+}$ *Bbs8*$^{fl/fl}$ and *Pdgfra*$^{cre/+}$ *Bbs8*$^{fl/fl}$ mice (n = 4).

### Figure S3

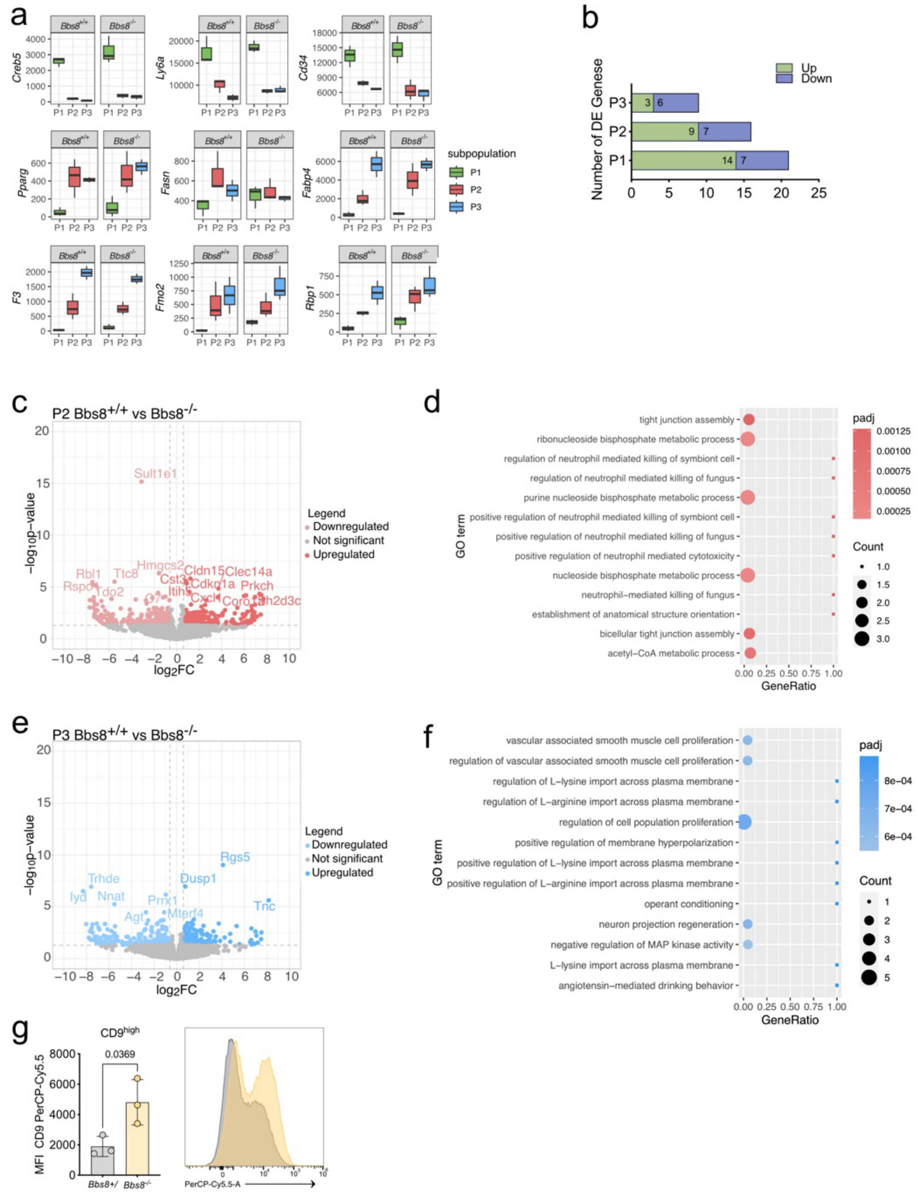


**Fig. S3. Phenotypic changes in APCs of lean *Bbs8*$^{-/-}$ mice. a**, Expression level of published P1, P2, and P3 markers split per genotype. Data are shown as mean ± SD. **b,** Differentially-expressed genes (DEG). Benjamini-Hochberg method was used to calculate multiple testing adjusted p-values. **c,** Volcano plots depicting the DEGs for P2 APCs. **d,** Over-representation analysis (ORA) of DEGs, highlighting the biological processes from gene ontology analysis in P2 *Bbs8*$^{-/-}$ compared to *Bbs8*$^{+/+}$ APCs. **e,** Volcano plots depicting the DEGs for P3 APCs. **f,** Over-representation analysis (ORA) of DEGs, highlighting the biological processes from gene ontology analysis in P3 *Bbs8*$^{-/-}$ compared to *Bbs8*$^{+/+}$ APCs. **g,** Median Fluorescent Intensity (MFI) of CD9-PerCP-Cy5.5 signal of PDGFR$\alpha^{+}$ cells from *Bbs8*$^{+/+}$ and *Bbs8*$^{-/-}$ mice (left). Histogram showing CD9-PerCP-Cy5.5 signal from concatenated *Bbs8*$^{+/+}$and *Bbs8*$^{-/-}$ files (right). Data are shown as mean ± SD, p-values have been determined using an unpaired Student’s t-test (n = 3).

**Figure S4**

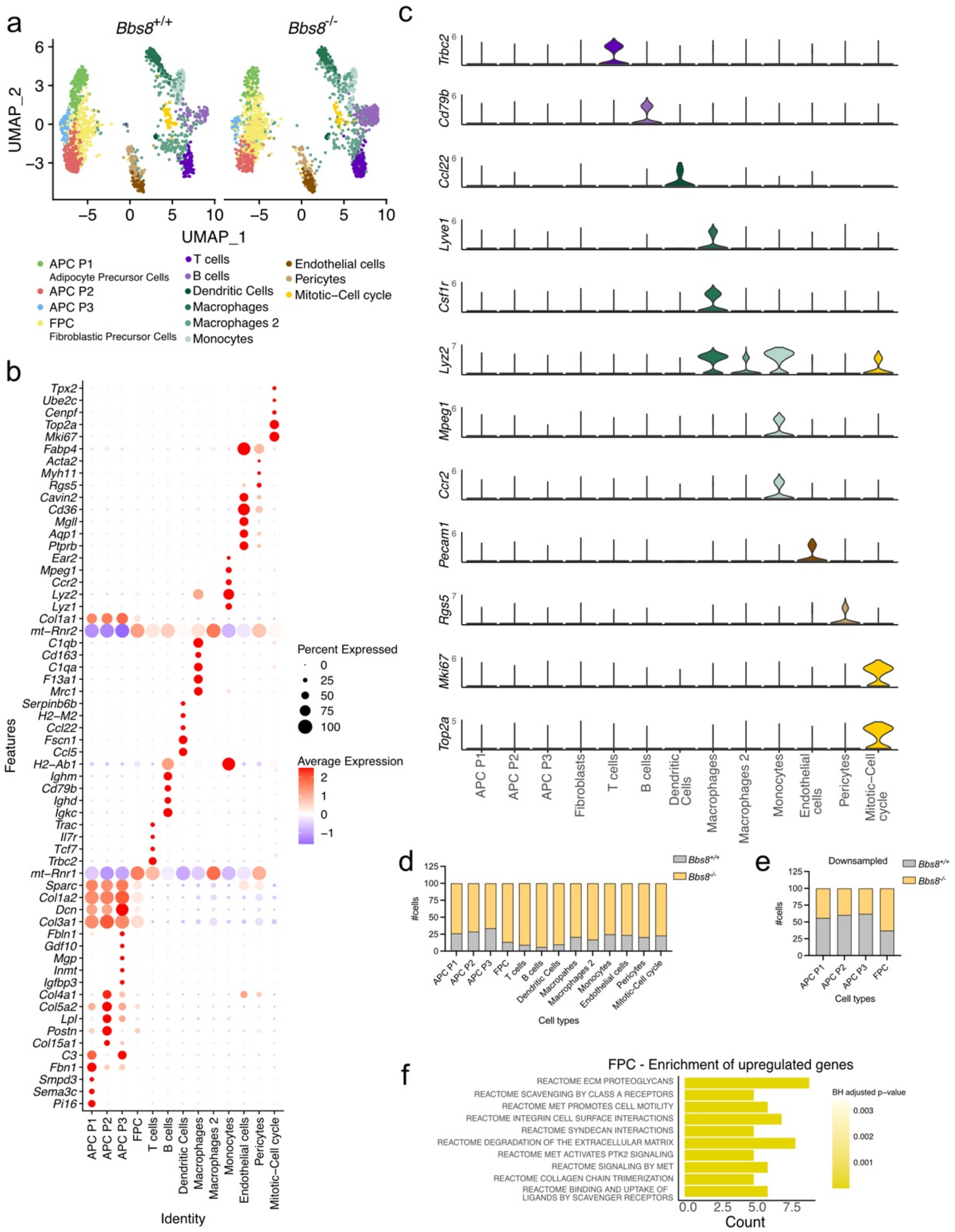


**Fig. S4. Loss of *BBS8* drives P1 cells into fibrogenic cells. a,** UMAP plot analysis on iWAT SVF from *Bbs8*$^{+/+}$ and *Bbs8*$^{-/-}$ mice split per genotype. **b,** Dot plot for top 5 marker genes in all cell clusters determined in Fig. S4a. **c,** Violin plots of published markers for all cell clusters determined in Fig. S4a**. d,** Relative cell numbers in each cluster per genotype. **e**, Relative cell numbers for the subsetted APC P1-P3 and FPC cluster. *Bbs8*$^{-/-}$ cells were down sampled. **f**, Reactome pathway analysis from genes upregulated in the FPC cluster.

### Supplementary Tables

**Tab. S1, DEGs of bulk RNA sequencing of P1, P2, and P3 from *Bbs8*$^{+/+}$ and *Bbs8*$^{-/-}$ mice (submitted as excel file)**

**Tab. S2, Single-cell identification marker for cell clusters (submitted as excel file)**

**Tab. S3, Sub-clustering of APC subpopulations (submitted as excel file)**

**Tab. S4, ORA_Reactome Enrichments FPCs (submitted as excel file)**

**Tab. S5, Flow cytometry antibodies**

| Antibody epitope | label | Producer | Cat. number |
|---|---|---|---|
| Streptavidin | BV785 | Biolegend | 405249 |
| CD31 | Biotin | Biolegend | 102503 |
| CD31 | BV785 | Biolegend | 102435 |
| CD45 | Biotin | Biolegend | 103103 |
| CD45 | AF700 | Biolegend | 147716 |
| TER119 | Biotin | Biolegend | 116203 |
| TER119 | AF700 | Biolegend | 116220 |
| CD26 (DPP-4) | PE/Cy7 | Biolegend | 137809 |
| CD9 | PerCP/Cyanine5.5 | Biolegend | 124818 |
| CD140a (Pdgfra) | BV421 | Biolegend | 135923 |
| CD55 | APC | Biolegend | 122513 |
| CD29 | PerCP-efluor710 | eBioscience (Thermo) | 46-0291 |
| CD142 | PE | SinoBiological | 50413-R001 |
| SCA1 | BV510 | Biolegend | 108129 |
| CD54 | APC-Fire 750 | Biolegend | 116125 |

**Tab. S6, Antibodies for immunocytochemical, fluorescence stainings**

| A) Primary antibodies | | | |
|---|---|---|---|
| **Antibody epitope** | **Species** | **Dilution** | **Cat. number** |
| Smo | Rabbit | 1:500 | Anderson Lab |
| Arl13b | Mouse | 1:1000 | Abcam ab136648 |
| Arl13b | Rabbit | 1:500 | Proteintech 17711-1-AP |
| Tubulin, gamma | Mouse | 1:2000 | Sigma T6793 |
| FABP4/A-FABP4 | Goat | 1:400 | R&D Systems AF1443 |
| B) Secondary antibodies | | | |
| **Antibody** | **Conjugate** | **Dilution** | **Cat. number** |
| Goat anti-mouse IgG1 (y1) | A488 | 1:2000 | Thermo Fischer #**A21121** |
| Donkey anti-rat | A488 | 1:500 | Dianova 712-545-153 |
| Donkey anti-rabbit | Cy3 | 1:250 | Dianova 711_165_152 |
| donkey anti-rat | A647 | 1:150 | Dianova 712-605-153 |
| Donkey anti-goat | A647 | 1:75 | Life Technologies A21447 |
| Goat anti-mouse IgG2a (y2a) | A647 | 1:1000 | Thermo Fischer #**A21241** |

**Tab. S7, Primer sequences**

| Gene name | Fwd sequence | Rev sequence |
|---|---|---|
| *Col1a1* | GAGATGATGGGGAAGCTGGC | CTCGGTGTCCCTTCATTCCG |
| *Col5a1* | CTTGTCCGATGGCAAGTGGC | CATCATCCAGAATCCGGGAGC |
| *Col6a1* | CAGGTACTACCGGTGTGACC | GAAGTACTTGACCGCATCCAC |
| *Loxl2* | CTGCCTGGAGGACACTGAGT | CGGTGATGTCTATCCACTGGC |
| *Fibronectin* | CTCCGAGACCAGTGCATCG | GAATCTTGGCACTGGTCAATGG |
| *Sparc* | CTGTGCCGAGAGTTCCCAG | CAGCAACTTCAGTCTGCTGAG |
| *Gli1* | TACCATGAGCCCTTCTTTAGGA | GCATCATTGAACCCCGAGTAG |
| *Ptch1* | GCCTTGGCTGTGGGATTAAAG | CTTCTCCTATCTTCTGACGGGT |
| *Tatabp* | GAGCTCTGGAATTGTACCGCAG | CATGATGACTGCAGCAAATCGC |
| *Gapdh* | AGGTCGGTGTGAACGGATTTG | TGTAGACCATGTAGTTGAGGTCA |
